## Supplemental tables and figures for "The interplay of DNA repair context with target sequence predictably biases Cas9-generated mutations"

### Supplementary Tables

| Knockout | Pathway | Function during DSB repair |
|---|---|---|
| Ercc1 | HR | Removes non-homologous 3′ single-stranded tails in HR and NHEJ. Also active in nucleotide excision repair and interstrand crosslink repair. [1] |
| Rad52 | HR | Promotes HR by displacing RPA from resected ends, allowing Rad51 to bind and start homology search.[2] |
| Trp53 | HR | Multifunctional tumour suppressor gene primarily inducing cell cycle arrest and apoptosis in response to DNA damage.[3] |
| Wrn | HR | Helicase and exonuclease, promotes NHEJ by suppressing resection.[4] |
| Lig1 | MMEJ | Ligates broken ends in MMEJ.[5] |
| Lig3 | MMEJ | Ligates broken ends in MMEJ. Also essential for mitochondrial DNA repair.[5] |
| Parp1 | MMEJ | Binds to resected DNA ends and biases pathway choice towards MMEJ.[6] |
| Nbn | MMEJ | Part of the MRN complex which performs resection of DSBs, initiating MMEJ or HR.[7] |
| Polq | MMEJ | Encodes Polymerase Theta, which displaces RPA from ssDNA, anneals sequences with microhomology and adds untemplated and templated nucleotides to the ends of DSBs.[8] |
| Dclre1c | NHEJ | Encodes Artemis, an endo- and exonuclease active during NHEJ.[9] |
| Poll | NHEJ | Polymerase. Prefers substrates with complementary sequence between ends. Also active in MMEJ.[10] |
| Polm | NHEJ | Polymerase. Prefers downstream base as template.[10] |
| Trex1 | NHEJ | A 3’ to 5’ exonuclease. Essential for ssDNA-mediated gene editing.[11] |
| Trp53bp1 | NHEJ | Protects broken ends from resection by MRN complex and helps recruit other repair factors.[12] |
| Lig4 | NHEJ | Primary ligase for NHEJ. Forms a synaptic complex with Xlf and Xrcc4 to hold broken ends together.[13] |
| Prkdc | NHEJ | Encodes DNA-PKcs, a kinase that binds to Ku complex and enables end processing by Artemis (product of Dclre1c). Regulates cell cycle arrest in response to DSBs.[14] |
| Xlf | NHEJ | Forms a synaptic complex with Lig4 and Xrcc5 to hold broken ends together.[13] |
| Xrcc5 | NHEJ | Forms a synaptic complex with Lig4 and Xlf to hold broken ends together. Stimulates activity of Lig4 for ligation of broken ends.[13] |

**Supplementary Table 1.** The panel of DNA repair gene knockouts used in this study, organised by pathway. Pathway assignments for each gene are based on the description of the function presented.

| Target | sgRNA | Forward primer for short-read sequencing | Reverse primer for short-read sequencing |
|---|---|---|---|
| HPRT exon 2 | TATACCTAATCATTATGCCG | GCAGATTAGCGATGATGAACC | ACACCACACACACACCCTCT |
| HPRT exon 3.1 | ACTTGCTCGAGATGTCATGA | TGTGTCCTGTAAAAGTTTAATGTGTAA | AATCCAGCAGGTCAGCAAAG |
| HPRT exon 3.2 | AGCCCCCCTTGAGCACACAG | GGACTGAAAGACTTGCTCGAGAT | CAGTAGCTCTTCAGTCTGATAAAA |
| HPRT exon 3.3 | GCCCTCTGTGTGCTCAAGGG | GGACTGAAAGACTTGCTCGAGAT | CAGTAGCTCTTCAGTCTGATAAAA |
| HPRT exon 3.4 | AAAATCTACAGTCATAGGAA | AAGTTCTTTGCTGACCTGCTG | TATGTAGCATAGTTTGCACAAGTT |
| HPRT exon 4 | GATGATCTCTCAACTTTAAC | CACAAGAATTTGCATTTAAGTGATG | CCTCTAAGTAAGTGGTTGAAAGC |
| HPRT exon 7 | TTAACAGCTTGCTGGTGAAA | GTAGTCTTTCCATATGCC | AACCCTAAGCCATTTCATAGT |

**Supplementary Table 2**. PCR primers used to amplify endogenously targeted sequences

### Supplementary Figures

**a**

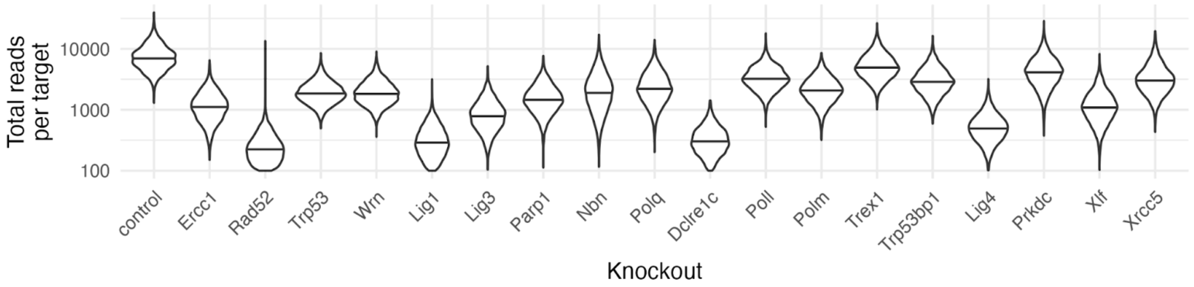


**b**

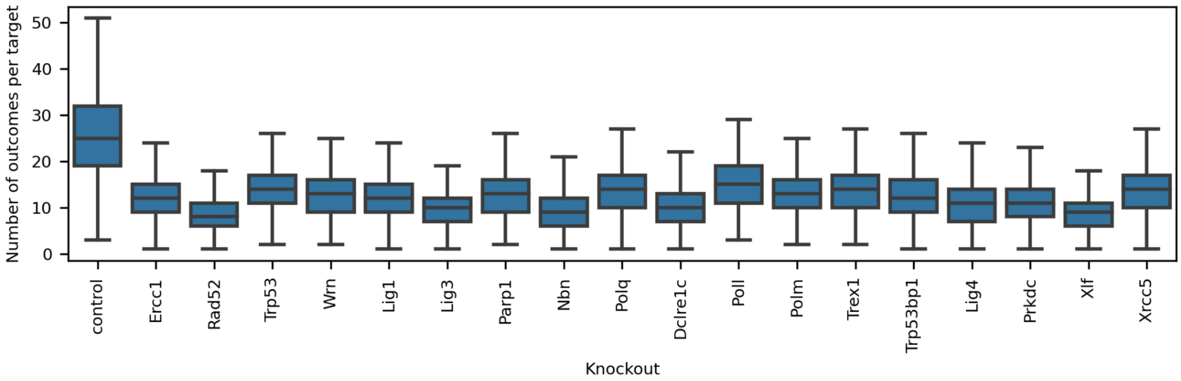


**c**

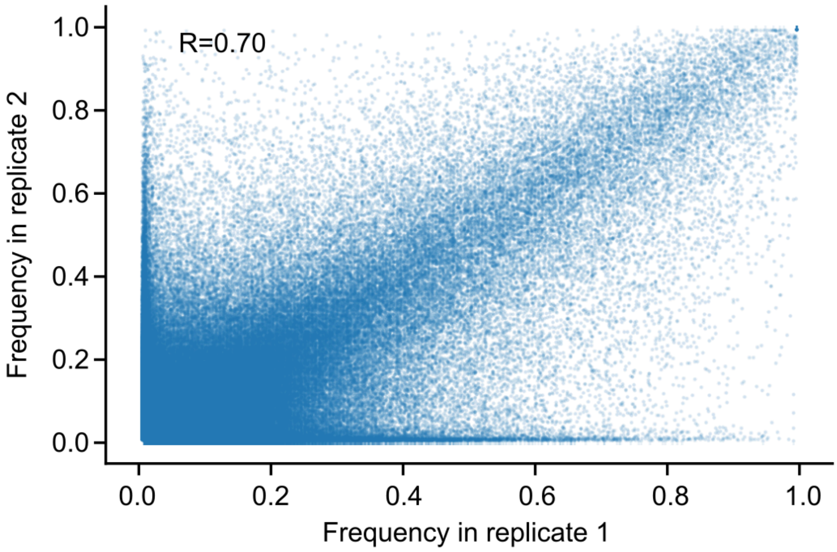

**d**

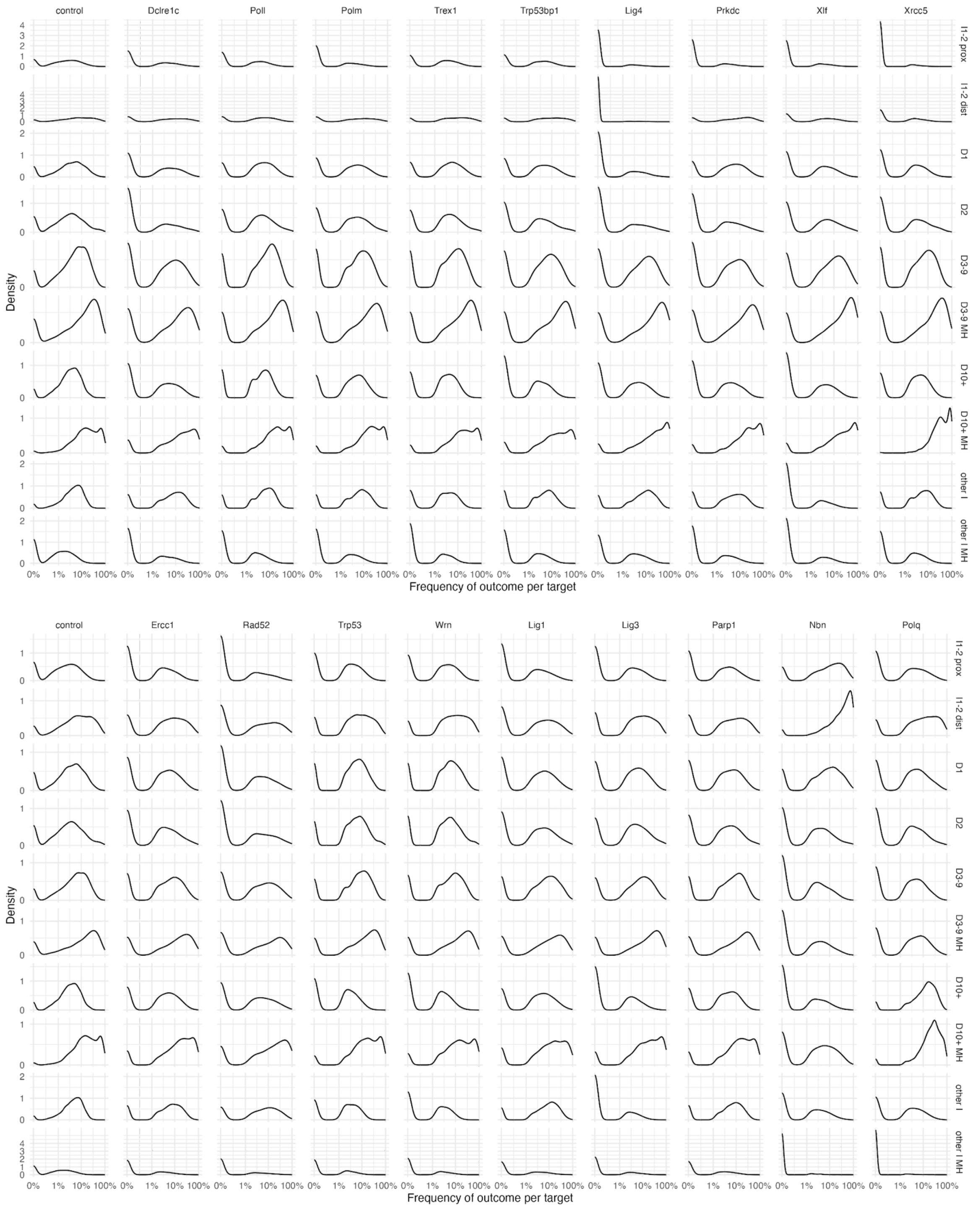


**Supplementary Figure 1**. **(a)**. Density (x-axis) of number of reads per target (y-axis; logarithmic scale) for screens in mouse embryonic stem cell lines with different repair gene knockouts (x-axis violins). Horizontal bar: median. Every target had at least 100 reads in every knockout. **(b)**. Number of outcomes per target (y-axis) for screens in mouse embryonic stem cell lines with different repair gene knockouts (x-axis). Box: median and quartiles; whiskers: 1.5x interquartile range. **(c).** Representative reproducibility of outcomes. Frequency of individual repair outcomes (markers) in replicate 1 (x-axis) and replicate 2 (y-axis) of one screen. Frequency of 1 = 100%. **(d)**. Diversity of outcomes. Frequency across targets (y-axis) of repair category outcome frequency within target (x-axis, logarithmic scale) of different outcome categories (rows) in different knockouts (columns).

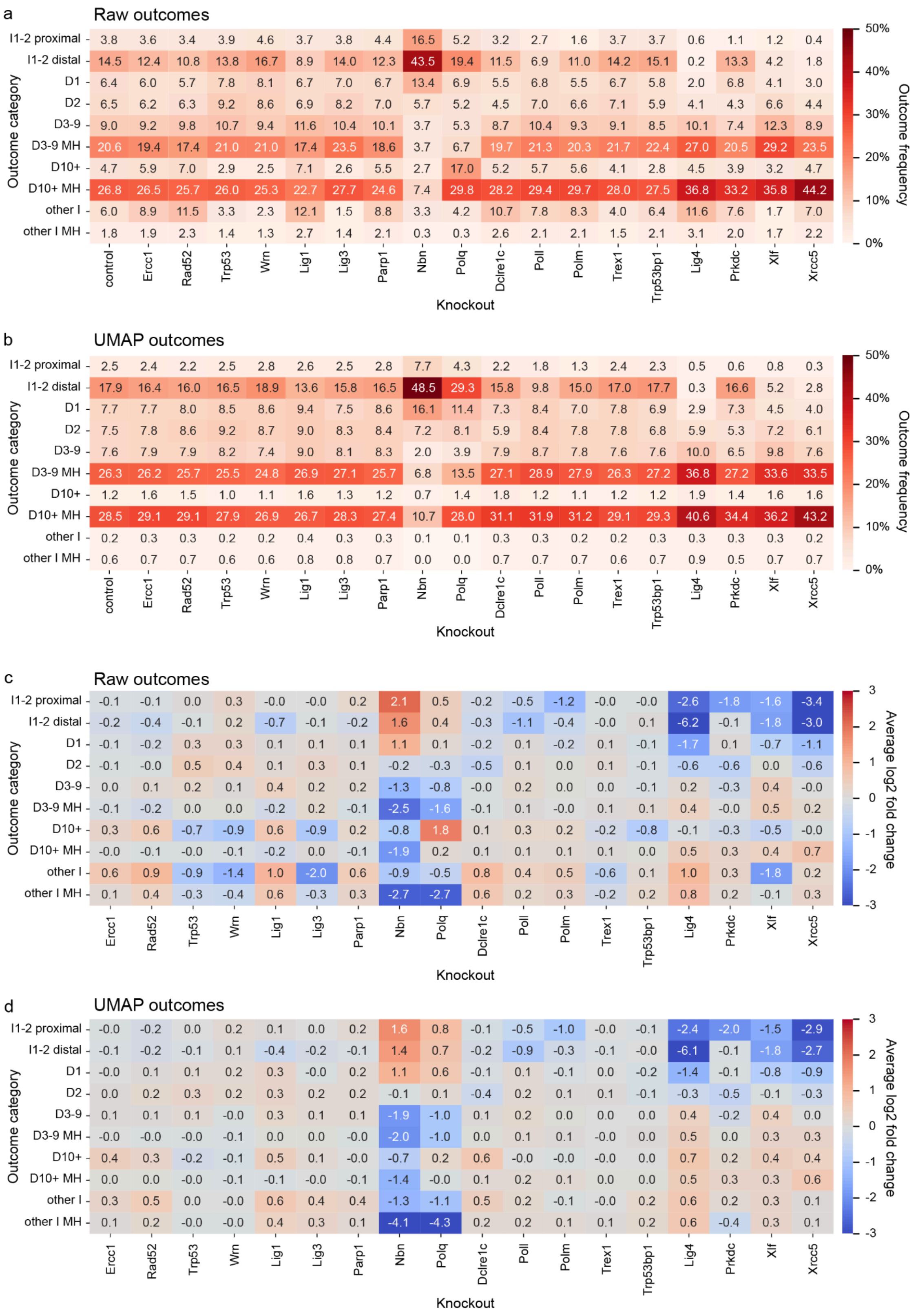


**Supplementary Figure 2. (a,b)** Average fraction of mutated reads across all targets (annotation, colour) of each outcome category (y-axis) observed in each knockout (x-axis). **(c,d)** Average of log-fold change across all targets (annotation, colour) of each outcome category (y-axis) observed in each knockout (x-axis). **(a,c)** Shows raw outcomes and **(b,d)** shows UMAP-filtered outcomes. D=deletion, I=insertion, MH=microhomology.

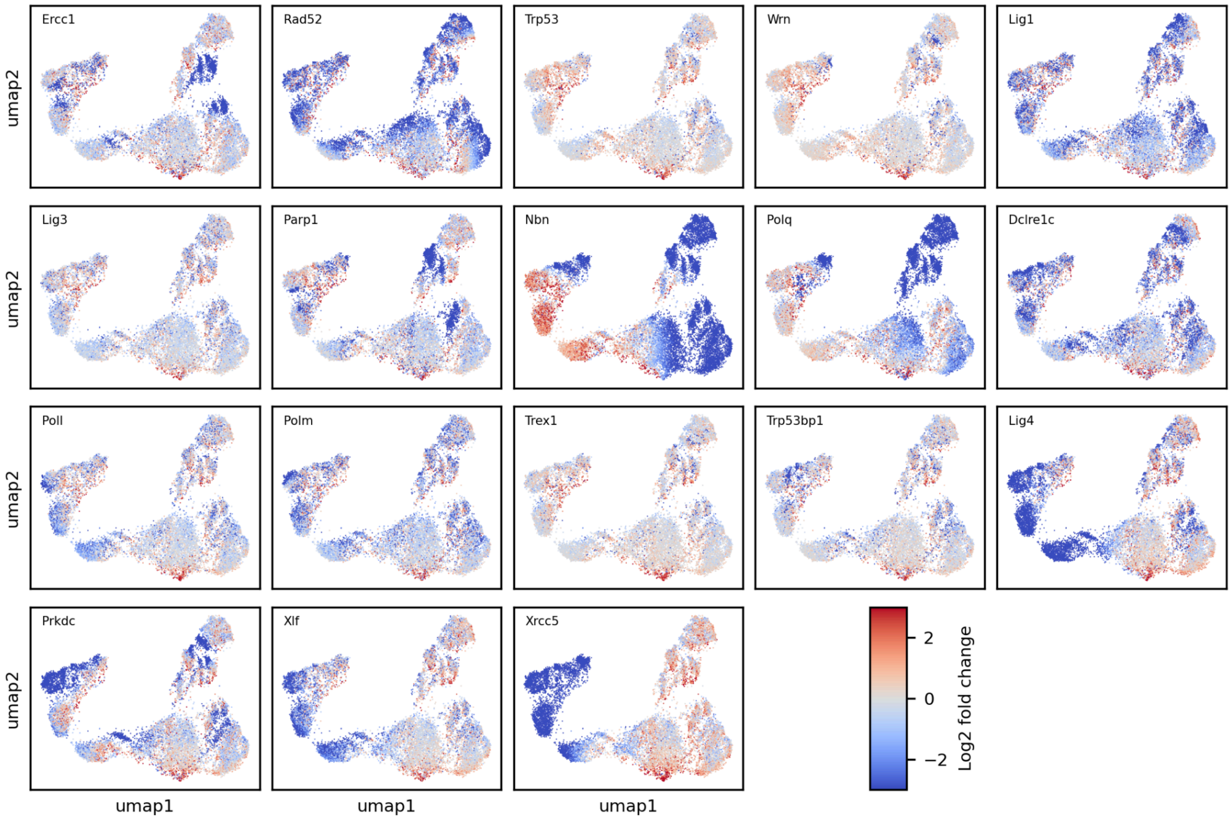


**Supplementary Figure 3.** UMAP projection of outcomes coloured by log-fold change in each knockout. UMAP projection of outcomes coloured by log-fold change (colour) of each outcome (dot) in each knockout (panels).

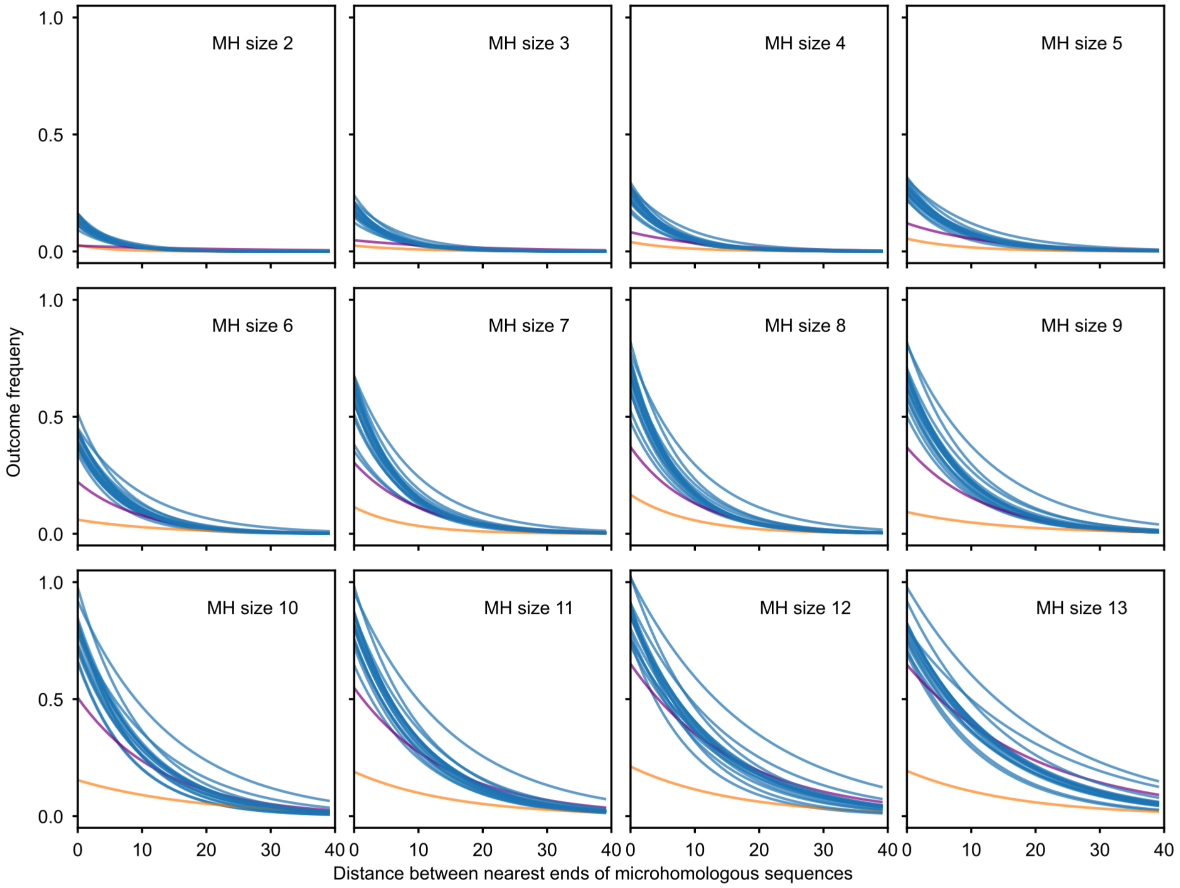


**Supplementary Figure 4.** The relationship between outcome frequency, size of microhomology sequence, and distance between microhomology sequences. Frequency of microhomology deletion (y-axis) vs the distance between microhomology sequence (x-axis) for a single length of microhomology (panels, labels) for each knockout (blue lines), with Nbn (orange) and Polq (purple) highlighted. Frequency of 1 = 100%.

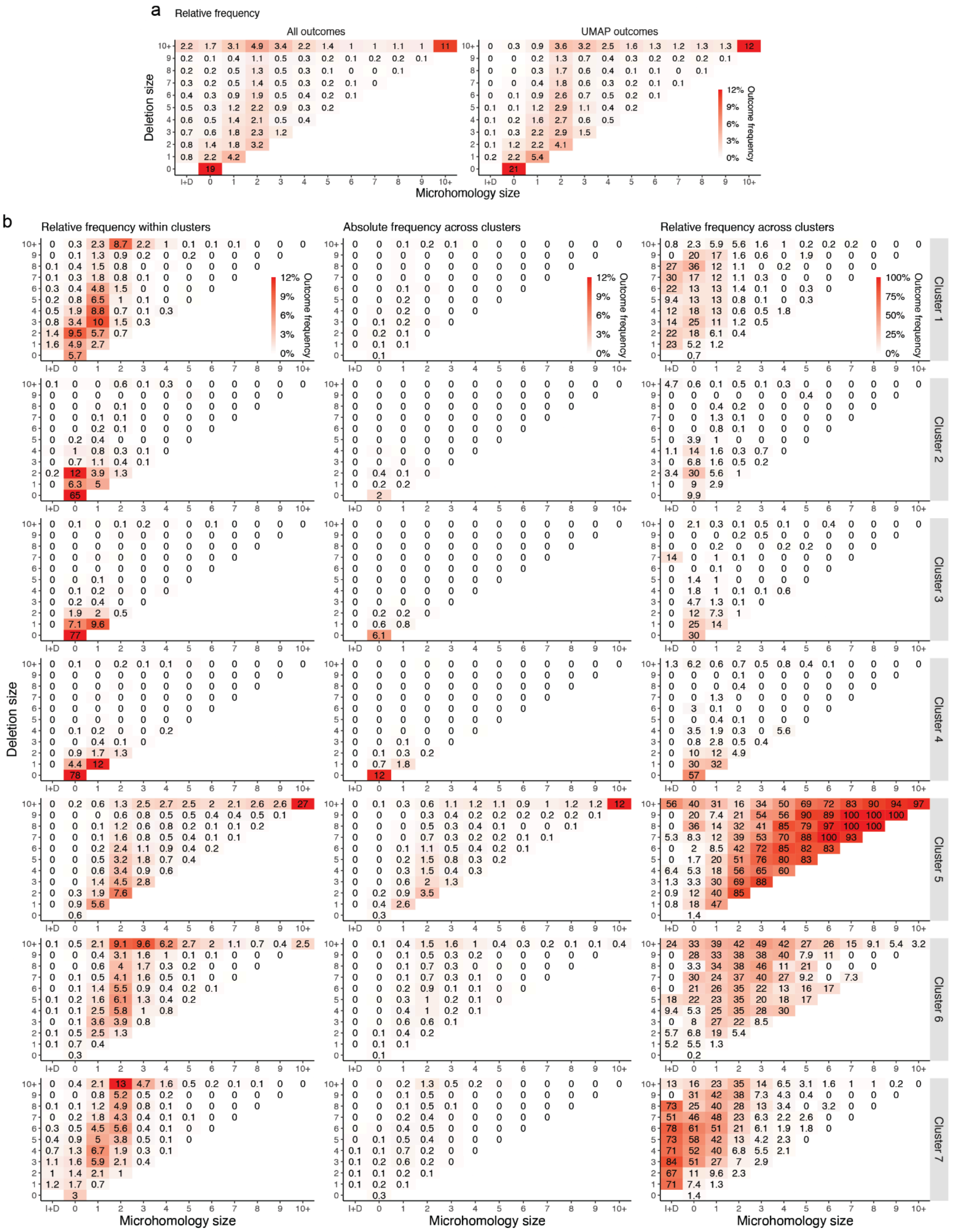


**Supplementary Figure 5.** Outcome distribution in terms of deletion size and microhomology. **(a)** Relative frequency of all outcomes or UMAP outcomes only. **(b)** Outcomes broken by UMAP cluster. Left column: relative frequency within cluster - each heatmap adds up to 100%. Middle column: absolute frequency distributed by cluster - same cell added up across clusters equals total UMAP frequency in (a). Right column: relative frequency distributed by cluster - same cell added up across clusters equals 100%. I+D = outcomes with insertions and deletions. Box at (0,0): pure insertions.

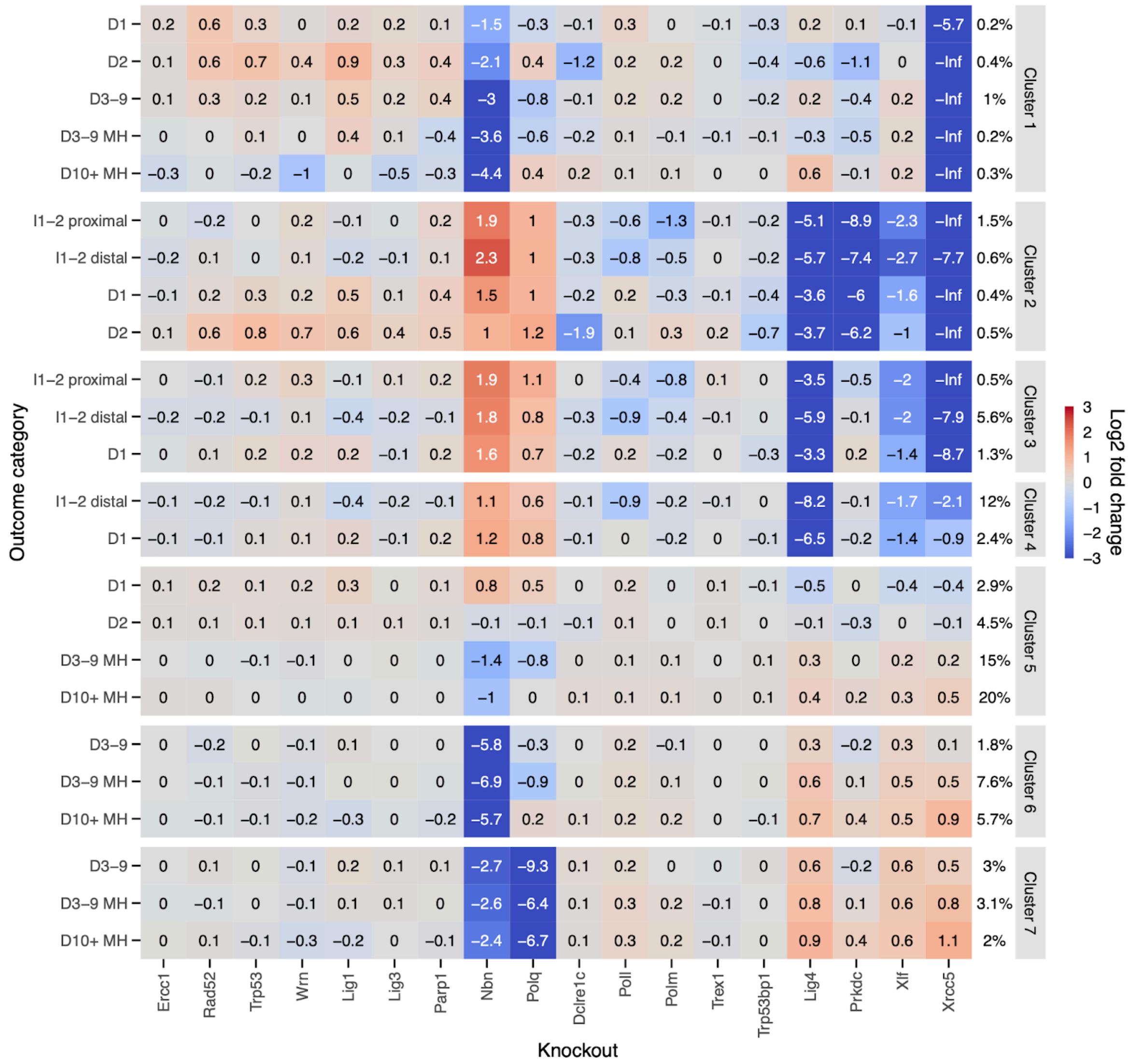


**Supplementary Figure 6.** Enrichment or depletion of outcomes categories within clusters across knockouts. Only outcome categories that made up >5% of the cluster depicted (a total of 7.2% outcomes not shown). D=deletion, I=insertion, MH=microhomology. Column next to cluster names indicates absolute average frequency of outcome category in a given cluster in control cells.

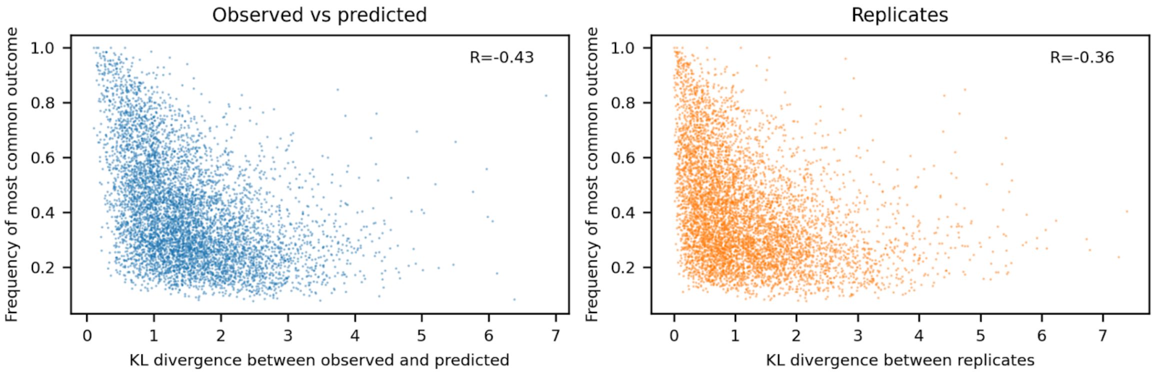


**Supplementary Figure 7.** Replicate concordance and prediction accuracy reflect frequency of most common outcome. KL divergence (x-axis) between observed and predicted outcome profiles (left panel, blue colour) or between replicates (right panel, orange colour) for one target (markers) contrasted against frequency of most common outcome for the target (y-axis). R = Pearson's R. Frequency of 1 = 100%.

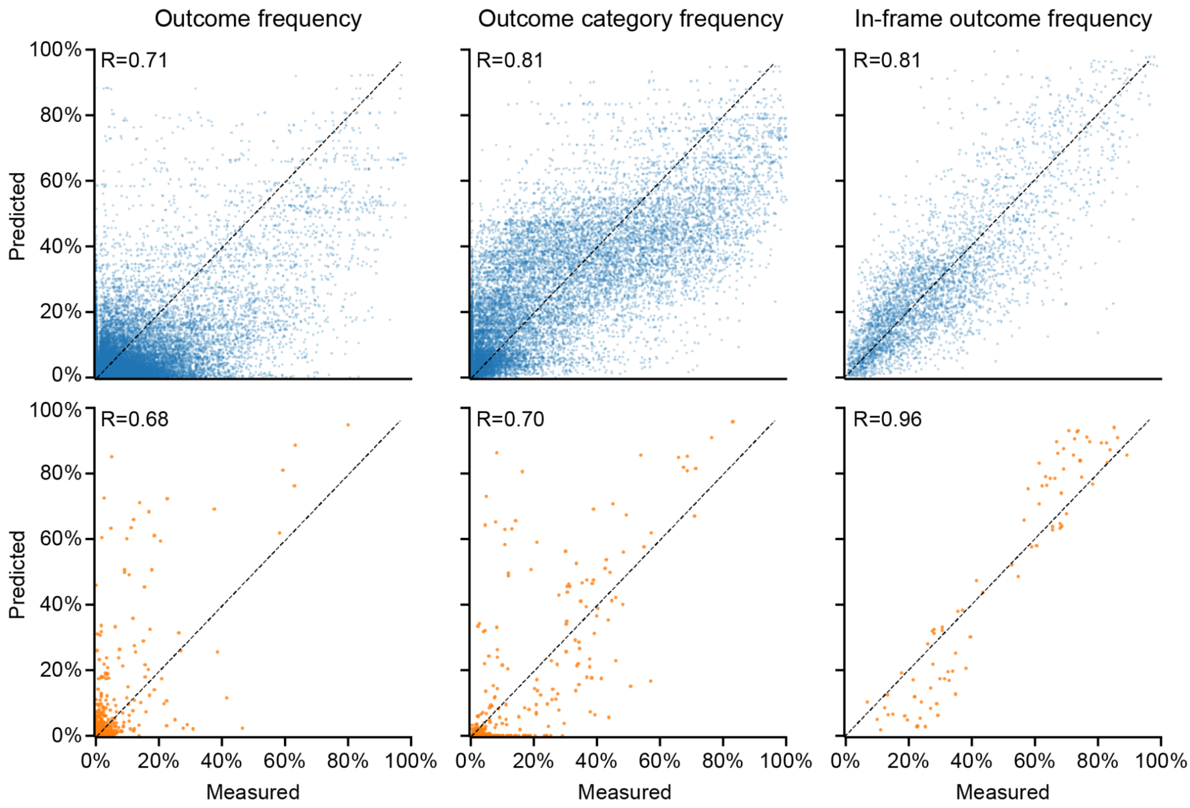


**Supplementary Figure 8**. Measured (x-axis) and predicted (y-axis) frequencies of individual outcomes, outcome categories and in-frame outcomes (columns) in held-out targets in Lig4, Polq, Poll, Polm, Xrcc5 knockouts. Top: original screen (N=670 targets), bottom: endogenous validation screen (N=7 targets). R = Pearson's R.
